# Single Cell Transcriptome Defines Cell Type Repertoire of Adult *Daphnia magna*

**DOI:** 10.1101/2024.05.29.596540

**Authors:** Indira Krishnan, Lev Y Yampolsky, Kseniya Petrova, Leonid Peshkin

## Abstract

Detailed knowledge of transcriptional responses to environmental and developmental cues is impossible without single cell (SC) resolution data. We performed two SC RNAseq experiments surveying transcriptional profiles of females and males of *D. magna*, a freshwater plankton crustacean *which* is both a classic and emerging new model for eco-physiology, toxicology, and evolutionary genomics. We were able to identify over 30 distinct cell types about half of which could be functionally annotated. First, we identified ovaries- and testis-related cell types by focusing on female- and male-specific clusters. Second, we compared markers between SC clusters and bulk RNAseq data on transcriptional profiles of early embryos, circulating hemocytes, midgut, heads (containing brain, eyes, muscles and hepatic caeca), antennae II, and carapace. Finally, we compared transcriptional profiles of Daphnia cell clusters with orthologous markers of 250+ cell types annotated in Drosophila cell atlas. This allowed us to recognize striated muscle cells, gut enterocytes, cuticular cells, as well as 5 different neuron types, including photoreceptors and 3 ovaries-related clusters, one of which tentatively identified as the germ line cells. One well-defined cluster showed a significant enrichment in markers of both hemocytes and fat body of *Drosophila*, but not with bulk RNAseq data from circulating hemocytes, allowing us to hypothesize the existence of non-circulating, fat body-associated population of hemocytes in *Daphnia*. On the other hand, the circulating hemocytes express numerous cuticular proteins suggesting their role, in addition to macrophagy, in wound repair. At the same time numerous cell types remain unidentified, including those that map to FCA groups ambiguously or are characterized by *Daphnia*-specific markers with no clear orthology in the fruitfly. Likewise, many known or presumed cell types or tissues in *Daphnia* have not been identified to SC clusters. A detailed in-situ hybridization study would be necessary to match not yet annotated SC clusters to functional cell groups.

**Highlights:**

- First single-cell transcriptomic atlas for *Daphnia magna*, identifies > 30 distinct cell types.
- Novel cell type representing circulating hemocytes may play a role in cuticle regeneration.
- Evidence for non-circulating hemocyte-like cells associated with the fat body in Daphnia.
- Cuticle/epithelial cells expressing photoreceptors, suggesting light-sensing capabilities.
- Subfunctionalization of divergent paralogs across cell types for ecological versatility.

## Introduction

Single cell (SC) transcriptomics is rapidly becoming the methodology of choice to study development, systems biology, and disease (Aldridge & Teichmann 2020). The unique advantages include the ability to detect changes in gene expression of healthy and abnormal cells reflecting tissue- and development stage-specific differences. As of now extensive SC RNAseq experiments have been done on a handful of model organisms, including humans, mice, *Xenopus* frog, zebrafish, *Drosophila*, and the nematode *Caenorhabditis elegans*, resulting in several comprehensive cell atlases for these models (Li et al. 2022; Liao et al. 2022; Ghaddar et al. 2023; Petrova et al. 2024) Liao. There is a significant interest in expanding SC transcriptomics in general and creating cell atlases for other models, particularly those proven to be useful and logistically permissive for ecological genomics, pharmacotoxicology and aging research. Each new organism does not simply provide a way to evaluate the existing body of work and plan new experiment using such atlas in a literal sense in a specific organism, but also enhance the value of data on other organism since contrasting the gene function and cell types on a tree of life elicits aspects not visible in isolated analyses. Tracing the gene origins and gene age based on the coding sequence has met with various difficulties (Capra et al. 2013), since genes have complex phylogenies, evolve at different rates and processes like subfunctionalization can only be identified in light of a function. Observing a gene in constellations with others in cell type atlases is one scalable way to gain insight into gene functions; observing concordant changes in cell-specificity of gene expression yields insights in mechanisms of complex processes, such as aging (Lu et al 2023) or brain functioning (Emani et al. 2024).

The freshwater plankton crustacean *Daphnia* is both a classic eco-physiological and an emerging new genomics model organism. A century or more ago several key biological concepts were either first discovered or studied using *Daphnia* as a model, including the ideas of germline-soma separation (Weismann 1983), cellular immunity and macrophagy (Metchnikoff 1901), phenotypic plasticity (Woltereck 1909), and caloric restriction effect on longevity (Ingle 1933). It is perhaps denotative that some of these discoveries relied on the investigation of or had a direct impact on the studies of histology and cell type biology in *Daphnia.* More recently, in addition to the crucial role *Daphnia* plays in toxicology (Tkaczyk et al. 2021), the newly available genomic tools have placed *Daphnia* on the frontline of functional and evolutionary genomics, allowing to ask questions spanning from molecular genetics and parasitology to population phylogenomics and ecosystems biology (Colbourne et al. 2011; Ebert 2011; Miner et al. 2012; Angst et al. 2022; Fields et al. 2022; Dexter et al 2023). As evidence is mounting that phenotype develop due to cell-level transcriptional responses to environmental cues (Aldridge & Teichmann 2020), it becomes clear that most of these research directions would benefit from a cell type atlas as a tool for single cell transcriptomics so far missing from the *Daphnia* genomic toolset.

Here we are reporting the first attempt to meet this need by conducting two single cell RNAseq experiments resulting in tentative identification of a number of cell types, including highly confidently identifiable myocytes, epithelial cells, certain neurons and midgut cells, as well as less confidently supported identification of reproductive tissues, fat body and hemocytes. By matching *Daphnia* cell clusters to well-characterized *Drosophila* cell types (Li et al. 2022) we attempt to identify cell types and tissues that are functionally and transcriptionally conserved across Pancrustacea. In addition, we report the analysis of cell type-specific transcription of duplicated genes present in the *Daphnia* genome. Genomes of *Daphnia* species contain a significant number of expanded gene families each consisting of 2-5 paralogs, many arranged in tandemly duplicated clusters. It has been long proposed (Colbourne et al 2011) that these duplications may have been retained via ecological subfunctionalization contributing to *Daphnia* ecological versatility, and that they indeed show divergent expression patterns proportional to amino acid sequence divergence. Data on cell type specific transcription will allow us to evaluate the possibility of “classic” subfunctionalization among paralogs by tissue-specific transcription. We have identified several cell types in *Daphnia* corresponding to key tissues and organs, largely based on statistically significant overlap in transcriptional markers with known *Drosophila* cell types.

Cross-species cell type matching is generally limited by several constraints and is particularly challenging when it comes to divergent species. *Daphnia* and *Drosophila* are sufficiently divergent for many cell type losses, gains, and transcriptional changes to be expected, yet a heavy investment has been made in the recent years into Drosophila single cell annotation which made it an attractive cross-reference species. While *Branchiopoda* is firmly supported as the sister group to the clade containing insect and their closest crustacean relatives, the remipedians (von Reumont et al. 2012; Lozano-Fernandez et al 2019), divergence between branchiopods and insects is probably over 500 Mya (Giribet & Edgecombe 2019; Montagna et al. 2019). Along with sequence divergence, tissue specificity may have also significantly diverged in many gene families. While some cell types, such as neurons and myocytes, are likely to be conserved in all arthropods or even all metazoans, others may be *Daphnia-* or dipteran-specific. This constitutes a challenge to cross-species cell type annotation, but also presents an opportunity to infer patterns of cell type evolution. We expect particularly large changes in tissues that evolved anew or were lost during transition from aquatic to terrestrial habitats, such as respiratory and excretory tissues.

## Materials and Methods

### *Daphnia* provenance and maintenance

*Daphnia magna* clone IL-MI-8 originating from an intermittent Mamilla pond (Jerusalem, Israel; 31° 46’ 40” N 35° 13’ 14” E) was obtained from *Daphnia* stock collection at Basel University, Switzerland, and has been maintained in the lab for over 5 years in modified ADaM reconstructed pond water (Klüttgen et al. 1994; https://evolution.unibas.ch/ebert/lab/adam.htm) at the density of 1 adult *Daphnia* per 20 mL. Food (*Scenedesmus* algae grown on COMBO algal medium, Kilham et al. 1998) was added daily at the concentration of 2E6 cells / *Daphnia* and neonates were removed and water changed every 3-5 days.

### Tissues dissociation and SC library preparation

Prior to tissue dissociation *Daphnia* were maintained for 3 days in antibiotics solution that contained 50mg/L of each of tetracycline HCl, streptomycin sulfate, and ampicillin sodium salt in 0.2 um-filtered ADaM medium, with the solution replaced daily. Five adult *Daphnia* (approx. 10 mg wet weight) were used for each SC transcriptomics sample. To disintegrate tissues with maximizing the yield of viable cells *Daphnia* were first placed in a beaker containing HBSS buffer without Ca^2+^ and Mg^2+^; 5.3 mOsm/L (ThermoFisher) on ice for 20 to 30 min to reduce tissues concentration of bivalent cations. After that *Daphnia* were transferred into enzyme mixture solution containing 8 mg/mL collagenase and 2.8 mg/mL (2000 U/mL) hyaluronidase (both SigmaAldrich) in HBSS dissociation buffer with 20mM sodium phosphate buffer, 25mM HEPES, 0.5 mM CaCl2 and 0.1 mg/mL BSA added; pH 7.2). Additionally, one of the samples in Experiment 1 was created by treating 15 individuals (approximately 30 mg weight weight) with collagenase only for the comparison of tissue disintegration efficiency. Eggs or embryos were removed from the brook chamber and *Daphnia* were cut transversely into three pieces with a microscalpel while in the enzyme mixture, with the rami of the antennae II removed (to prevent serrated setae present on the rami from clogging the microfluidics equipment). The samples were then placed in a shaker at 50 rpm, at 30 °C for 1 h. At the end of the incubation, tubes were placed on ice until big debris settled to the bottom and the cell suspension was passed through stacked 70 um & 30 um filters (pre-wetted with the dissociation buffer), and rinsed with 3 mL of the same buffer. The cell suspension was spun at 500 x g for 25 min at 4 °C. Supernatant was removed and the pellet was washed three times with the wash buffer (HBSS w/o Ca^2+^ and Mg^2+^, with 25mM HEPES added, pH 7.2) and suspended in 2 ml of the same buffer. Cell viability was tested by Calcein-AM/Ethidium homodimer III assay (Promokine). Briefly, working solutions of Calcein AM (400 uM) and Ethidium homodimer III (100 uM) were freshly prepared from 4 mM and 2 mM stocks, respectively, in the wash buffer. Four uL of each of the working solutions were added to 200 ul of cell suspension to a final concentration of 8 uM (Calcein AM) and 2 uM (EthD-III) and the mix was incubated in the dark at room temperature for 30 min. Controls without Calcein AM and EthD-III were included for each assay. Following the incubation, the cells were pelleted by centrifugation at 200 x g for 10 min and washed twice the wash buffer. The cells were then suspended in 100 ul of the same buffer and analyzed by flow cytometry on Moxi Go II flow cytometer, with calcein fluorescence (Ex/Em: 488 nm / 525 nm) indicative of live cells, and EthD-III fluorescence (Ex/Em: 488 nm / 650 nm) indicative of dead cells. Percent of cells in the Calcein AM+ quadrant was used as percent viability of the cell suspension. 10 ul of the original cell suspension was mixed with 1ul of 1mM Hoechst 33342 and cells counted under microscope in the DAPI channel to determine the number of cells/mL. Cells were then diluted to approximately 90,000 cells / mL in the wash buffer with 15% (v/v) of Optiprep before loading for transcriptome barcoding of single cells on the inDrop platform.

### Data analysis

The sequencing data was demultiplexed using the inDrops pipeline (https://github.com/indrops/indrops). Reads were filtered to remove those with low quality or low complexity and subsequently mapped to the repeat-masked *Daphnia magna* DM3 genome assembly (BioProject ID: PRJNA624896; Fields et al., personal communication) using STAR aligner (Dobin et al. 2013). The count matrices were further filtered to remove degraded cells (>15% of mitochondrial content). Drosophila and human orthologs for Daphnia genes were obtained using reciprocal best hit comparisons based on HMMER3 and BLOSUM45 amino acid substitution matrix reflecting evolutionary distance between these species (Menlove 2009).

### Visualization and clustering

In order to visualize and interrogate the data as well as provide an instant access to the community we used SPRING (Weinreb et al., 2018) for interactive analysis and visualization as well as to provide instant access to the community. All SPRING plots (k-NN graphs rendered using a force-directed layout) were generated using the default parameters in the SPRING upload server: https://kleintools.hms.harvard.edu/tools/spring.html. Clustering the data using default parameters in SPRING via the Louvain clustering method. Clusters identified by Louvain method will be hereafter referred to as L1, L2, etc.

### Cluster annotation

To functionally annotate clusters discovered by Spring tool we used our own single cell and bulk RNAseq data, data on published *Daphnia* tissue specific markers (limited to fat body (Goldmann et al. 1999) and ovaries (Toyota et al. 2017) and data on cell-type specific markers in the nearest relative with a well-characterized single cell transcriptomics, *Drosophila melanogaster*. Firstly, we contrasted, using Spring counter-select tool, female and male-derived libraries to narrow down on female- and male-specific cell types, primarily ovaries and testes. One should keep in mind that the female and male germlines may or may not be differentiated this way. Additionally, male-derived libraries may be enriched in cells specific to antennae I - structures that are much more developed in males than in females. These may or may not include gustatory, olfactory, and tactile receptors, but also might include antennae I-specific muscles. Similarly, we contrasted head- and body-derived libraries within female samples, expecting CNS and eye-specific transcripts to be overexpressed in the head libraries relative to the body libraries.

Second, we utilized the results of a *Daphnia* “gross anatomy” bulk RNAseq experiment (Dua and Yampolsky 2024), in which transcriptional profiles have been obtained for early stage embryos (<16 h after egg-laying; corresponding to stages 1-2 (Mittmann et al. 2014;Toyota et al 2016) and several body parts and tissues of their mothers. These included (in the order they were sampled from the specimen): hemocytes (the subpopulation circulating the hemolymph), midgut, antennae II, heads, carapaces and, finally, carcasses left after the removal of all the listed tissues. Transcripts abundance of each tissue was measured by Nanopore RNAseq experiment and compared to that in the carcass using DESeq R package, recording both transcripts over- and underrepresented in each body part. With the exception of hemocytes and, possibly, early embryos, all the body parts contain a variety of tissues and cell types. Specifically, we expect the gut to be enriched in enterocytes and smooth muscle cells, antennae II to be enriched in epithelium cells, striated muscle cells and neurons, carapace in epithelium cells, and heads in CNS neurons, eye photoreceptors, epithelium cells, striated muscles (because a large portion of swimming antenna-driving muscles is retained within the heads during head removal), and gut epithelium (because the hepatic caeca, the cul-de-sac gut appendages with presumed digestive enzymes secretion function (Smirnov 2017) are located within the head and are retained within it). On the other hand, the carcass is expected to be enriched, relative to each individual body part, in cell types associated with fat body, ovaries, nephridium and epipodites (sections of thoracal appendages responsible for osmoregulation, Smirnov 2017). It is not known whether the locomotory muscles (present in antenna II and the head) and gut muscles differ transcriptomically from the heart muscles and muscles driving filtering appendages (both present in the carcass), so there we had no *a priori* prediction about relative abundance of muscle-specific transcripts between these parts of the anatomy. On the other hand, carapace and hemocytes are expected to be depauperate in transcripts associated with any type of muscle. Finally, early ovaries are expected to be similar to the germline and possibly some ovary-associated cell types.

In order to match cell types’ markers to those of *Drosophila* cell types annotated in Fly Cell Atlas (hereafter FCA, Li et al. 2022) a subset of *Daphnia* genes with a recognizable *Drosophila* ortholog among the top 100 markers of each cell type was matched to cell type markers in FCA data. Two different criteria of inclusion of FCA markers were used. The less stringent cut-off included all genes with IsMarker=1 flag in FCA (corresponding to cell types relative enrichment of logFC>0 and adjusted p-value<0.05. The more stringent cut off was logFC>2. The results were fundamentally similar. The significance of the number of unique genes in this intersection was then determined using Fisher exact test (FET) using the number of genes in the marker set for each *Daphnia* and *Drosophila* cell types as a reference. Specifically, the 2×2 contingency table for each combination of observed *Daphnia* cell types and *Drosophila* cell types (FCA “groups”) consisted of the number of genes in the intersect between *Daphnia* and *Drosophila* markers list, the number of genes in the *Daphnia* cell type marker list outside of the intersect, the number of genes in the *Drosophila* cell type marker list outside of the intersect, and the number of unique genes in the union of all markers of all cell types in both species. The FET therefore tested, for each cell type in our data, the non-randomness of the intersect between *Drosophila* and *Daphnia* markers. The resulting FET p-values were adjusted for multiple testing by false discovery rate (FDR) with the total number of tests performed set to the product of the number of *Daphnia* cell types (38 in Experiment 1 dataset) and number of *Drosophila* cell types in FCA (244). This product was used as the total number of tests for the sake of being conservative, even though many of *Daphnia* and *Drosophila* cell types combinations resulted in 0 overlap, in which no test was conducted.

Finally, at least four clusters (three of them being cuticle cells) were annotated despite the lack of consistent enrichment with FCA markers based entirely on ontology of top few markers, without an attempt to test for non-randomness of functional enrichment. Ten out of 38 Louvain clusters identified by Spring (Experiment 1) were deemed artifactual and excluded from further analysis (but they were included into the calculation of FRD in enrichment analysis for the purpose of being conservative). These clusters were excluded based on the following criteria: 1) basal spring location either to all clusters, or to another, better defined clusters; 2) lack of high specificity markers (top markers with enrichment score Z<2); 3) high level of expression of ubiquitously expressed ribosomal and mitochondrial proteins; and 4) no clear matches between experiments 1 and 2. Additionally, we excluded from further consideration three clusters represented by less than 10 cells.

### Paralogs expression analysis

*Daphnia* paralogs were identified in two ways. First, in conjunction with the widespread use of *Drosophila* orthologs for cell type annotation, we analyzed *Daphnia* genes that were identified as orthologs to the same *Drosophila* gene, regardless of the degree of divergence between them. This approach is biased towards conserved proteins as it eliminates *Daphnia-* specific genes with no identifiable *Drosophila* ortholog from the analysis. Second, we did a blast search of *D. magna* proteome (DM3 annotation, the longest isoform used), against itself and retained pairs of paralogs (e<0.0001) in which areas of homology covered at least 50% of sequence length in at least one of the paralogs. To correlate transcriptional specialization with sequence divergence we calculated, for each pair of paralogs, transcriptional distance as euclidean distance in the space of 38 Louvain clusters using the difference between the two paralogs’ Z-scores in each cluster. This measure was calculated for all genes within top 1000 showing expression within each cluster, the default cut-off in Spring. If the other gene in the pair was not present among these 1000, its Z-score was set to 0.

## Results

### Tissue dissociation efficiency

The two versions of tissue dissociation protocol yielded similar results. Treating samples with the mix of collagenase and hyaluronidase resulted in a better cell count than with collagenase alone (5E4 vs. 1E4 cells per ml in the final suspension prior to pelleting). However, sequencing libraries prepared from samples treated in both ways resulted in similar barcode counts (3339 - 3418 cell per library) and similar representation of libraries in cell types (Fig.1 insert), indicating that the two treatments resulted in similar counts of viable cells and in homogeneous distribution of cell-type counts (Likelihood ratio Chi2 = 42.6, P>0.24, Supplementary Table S1), indicating that most tissue were disintegrated equally well.

**Figure 1.**
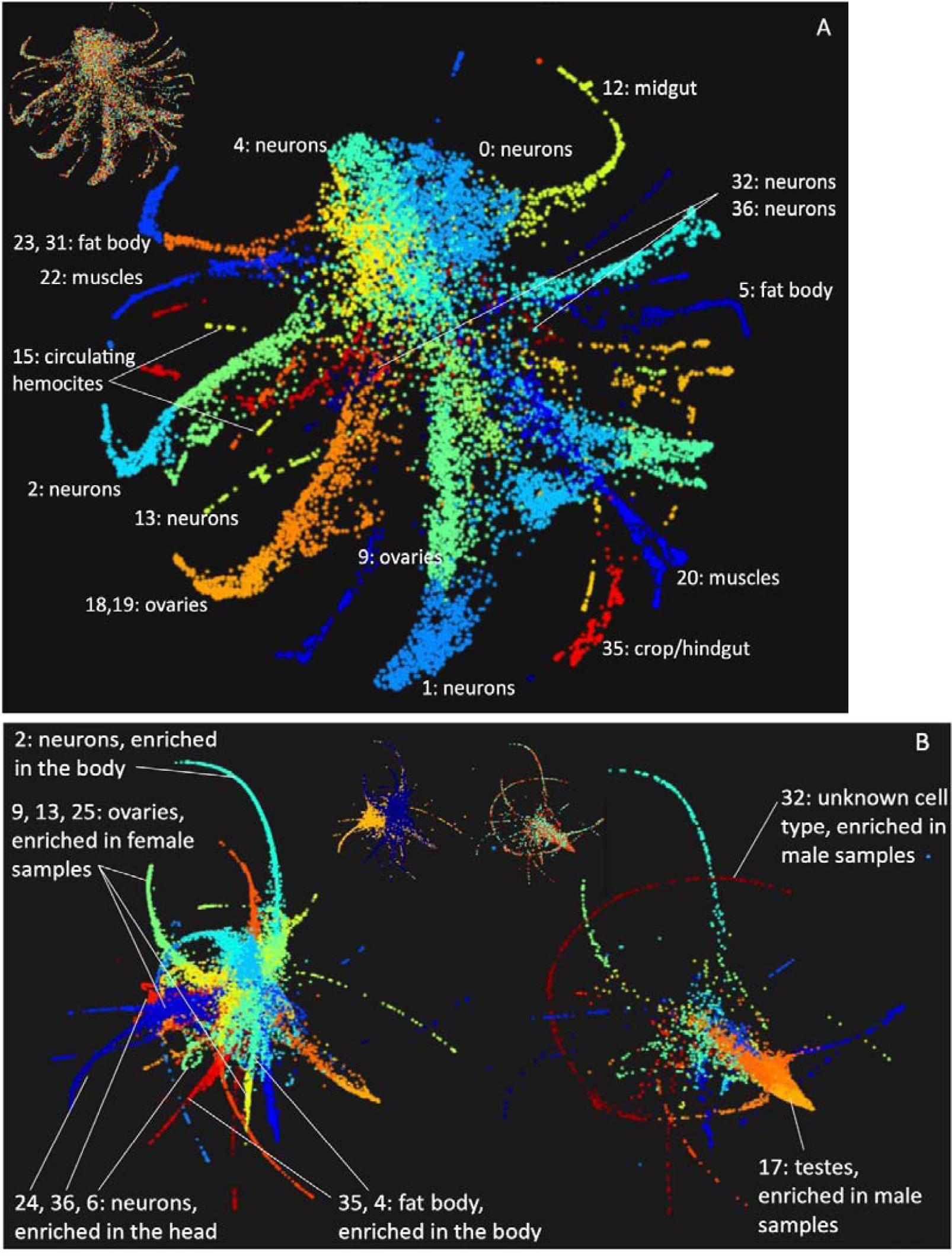
Results of two single cell transcriptomics experiments visualized by SPRING tool. Main panels: colors represent Louvain clusters; inserts: colors represent replicate or body-part specific libraries. A: Experiment 1 (available at https://shorturl.at/opAIV). Tentatively identified cell clusters (Table 1) labeled. B: Experiment 2 (https://shorturl.at/oDFM7). Female- (left) and male-derived libraries (right) shown separately. Insert: in female-derived libraries colors represent body parts (blue: bodies; yellow: heads); in male-derived libraries colors represent replicate libraries. Select clusters enriched in males vs. females or heads vs. bodies (Table 1) labeled. See Table 1 for all clusters overrepresented in sex- or body part-specific libraries.

**Table 1.**
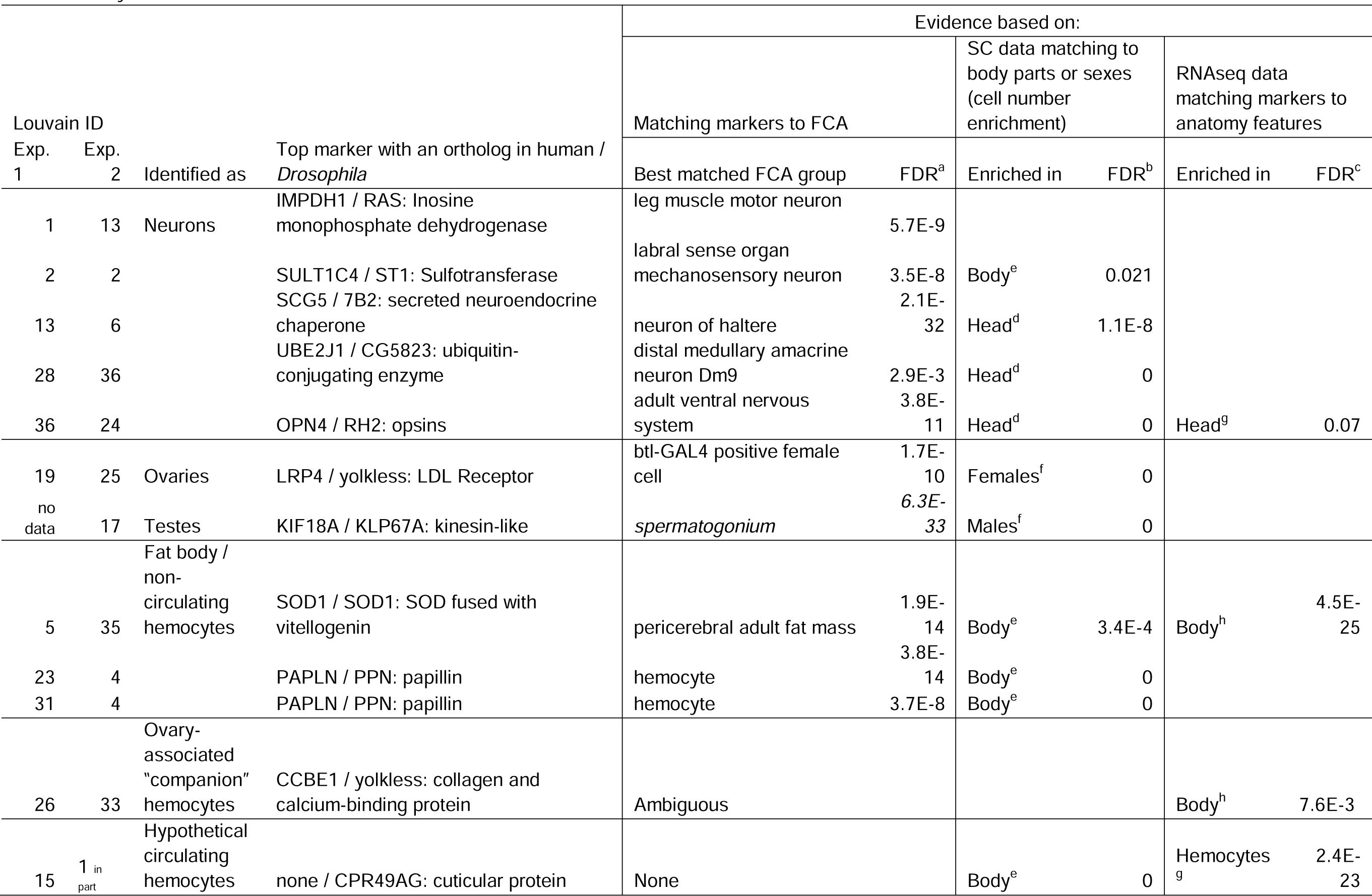
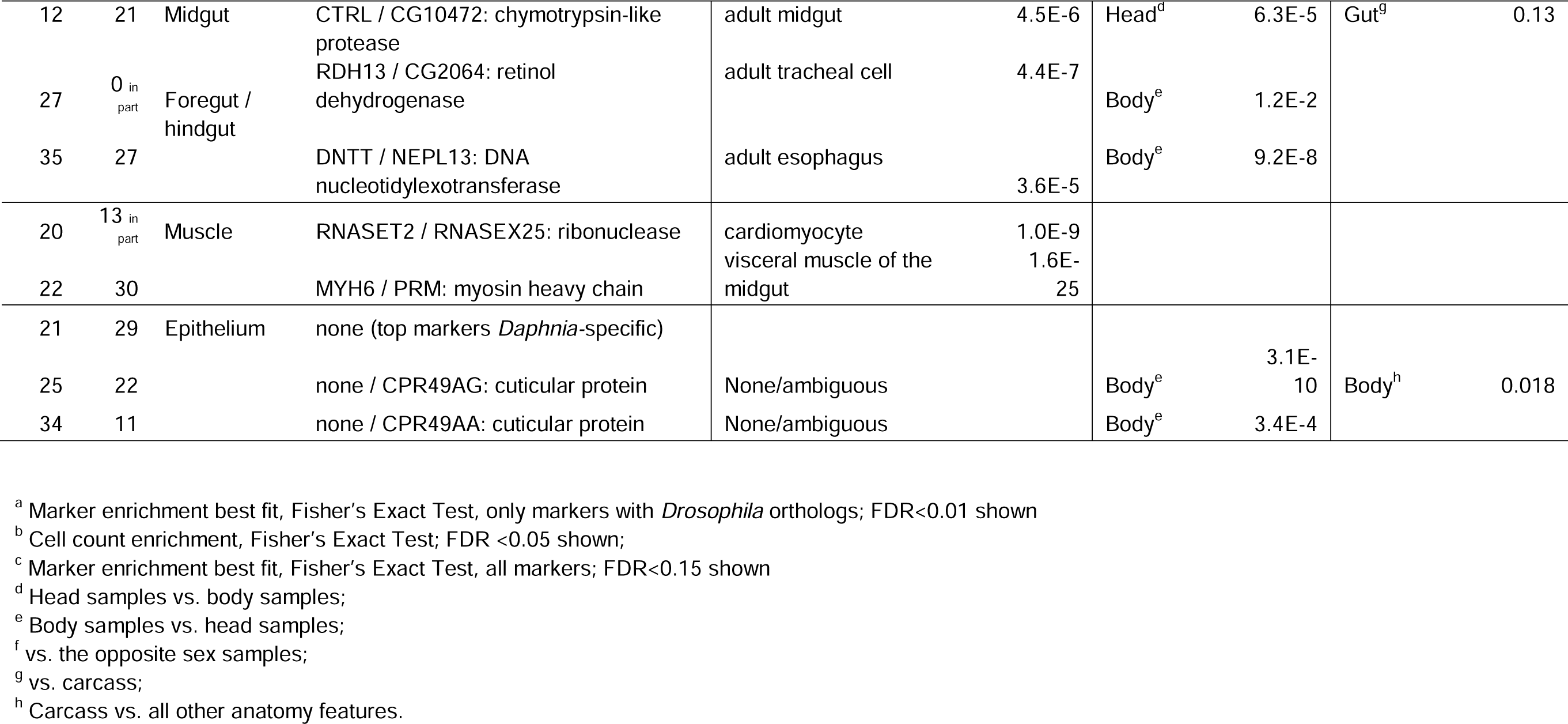
Cell clusters revealed in 2 single-cell RNAseq experiments (Exp.1 and Exp.2) tentatively identified by matching markers to Fly Cell Atlas cell types, sexes and anatomy features.

### Cell types annotation

In total, we have identified over 30 single cell clusters with distinct transcriptional profiles (Figs. 1, 2; Table 1). In the next sections we attribute and discuss these cell types which were identifiable either based on transcriptional similarity to *Drosophila* cell types (Li et al. 2022; Supplementary Table S2) or by matching to *Daphnia* body part-specific transcriptional profile. Ten clusters without strong evidence of transcriptional specificity are listed in Supplementary Table S3.

#### Neurons

Neural tissues in *Daphnia* occur throughout the body, with three morphological features being distinct in the head, namely the brain (or cerebral ganglion; this structure also includes the ocellus), the eye, and the optical lobe. Five cell clusters showed a highly significant enrichment with homologs of various types of marker genes originally defined in Drosophila neurons (Table 1; Supplementary Table S2, Fig 2, Supplementary Fig 1. Only one of these five hypothetical neuron types could be convincingly identified to a particular functionality, a small cluster L36 in which 4 out the top 5 and 5 out of 5 top markers in Experiments 1 and 2, respectively, were opsins, matching this cell type to photoreceptors. Furthermore, the only other identifiable marker of this cell type (in Experiment 2) was a homolog of *Drosophila* myosin ATPase *ninaC*, a protein required for photoreceptor cell function. Not surprisingly, this cell type showed a strong enrichment in the head-derived samples. There was an additional cluster (L25, Fig. 1) which showed a mild enrichment in markers of photoreceptor cells and antenna glial cells (Supplementary Table S2), but with no indication of opsins or any other receptor-specific proteins being differentially expressed. Instead, this cluster overexpresses several homologs of *Drosophila* cuticular proteins, possibly linking it to epithelium cells (see below) with shared markers with receptor cells being an artifact.

**Figure 2.**
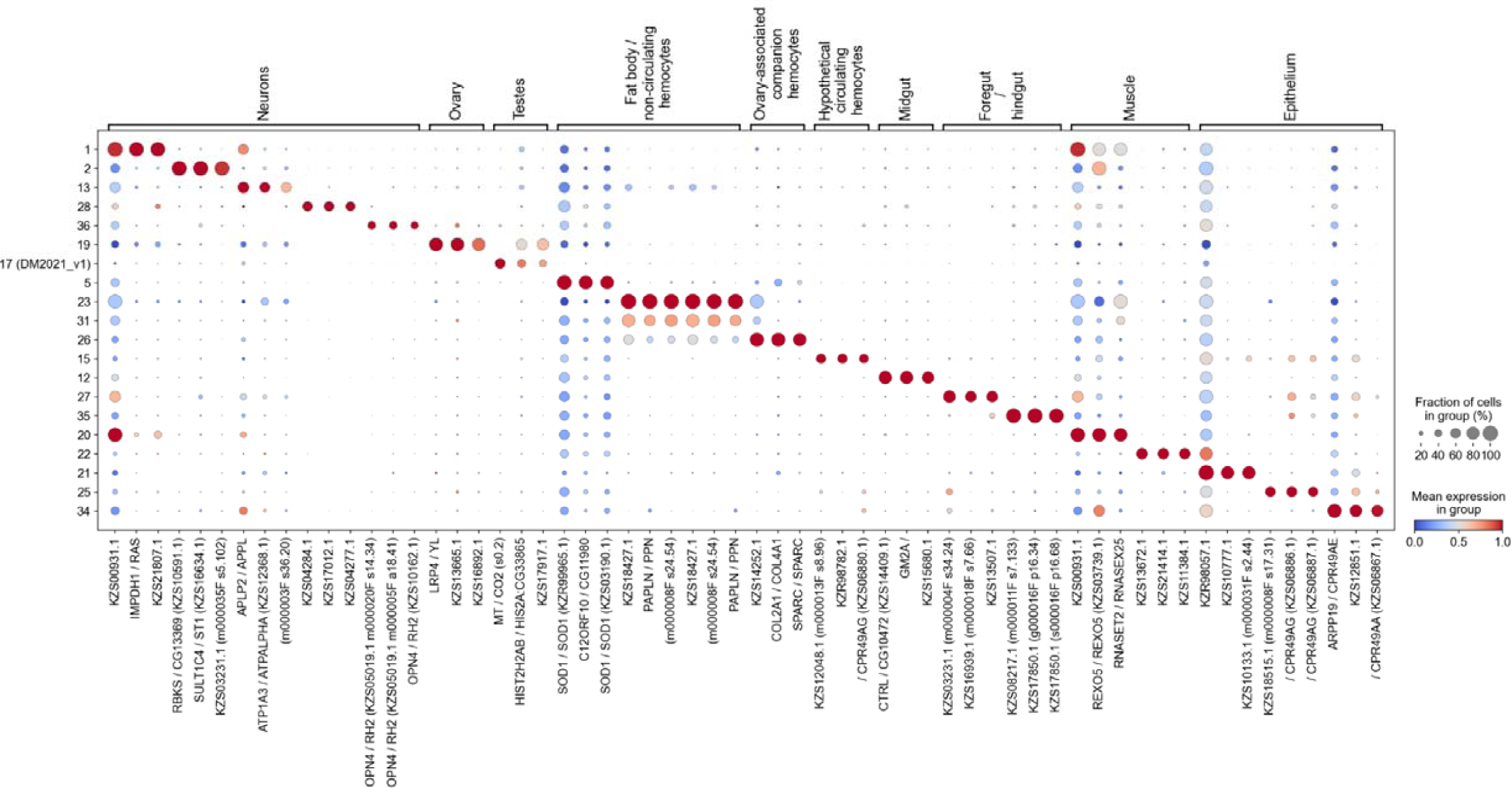
Dotplot of several identifiable cell types showing fraction of cells with at least one read and mean expression in each cell type. See Supplementary Fig.1 for more detailed dotplots for reproductive and neural tissues cell types

Other neuron-like cell clusters were harder to identify to a particular neuronal type. Three of these also showed enrichment in the head vs. body (L13, L28, L36), while L2 was enriched in the body (Table 1). With little concordance with these enrichment associations, these clusters also showed significant matches to a variety of *Drosophila* neuron types, including adult ventral nervous system, leg muscle motor neurons, mechanosensory neurons, among others. Markers overexpressed in one or more of these cell types that match FCA markers and markers identified as neuronal by Croset et 2018 and Cho et al 2020 included, besides the aforementioned opsins, the homologs of *Drosophila amontillado (PCSK2 / AMON), bruno-3 (CELF6 / BRU3), Galphao (GNAO1 / GALPHAO), hamlet (PRDM16 / HAM), rab3 (RAB3C / RAB3), rhoGAP (SYDE2 / RHOGAP100F), as well* one of several *Daphnia* paralogs of apolipoprotein *neural lazarillo (APOD / NLAZ*). Note that the majority of these markers were strongly specific to only one of the five presumed neuron clusters.

#### Myocytes

Muscle tissues occur nearly throughout the body of *Daphnia* (Smirnov 2017 Fig. 1.3), including the head where locomotory antenna driving muscles are located, antennae themselves, thorax, where muscles drive filtering and respiratory appendages, abdomen, gut, and heart. Myocytes were among the most reliably identified cell types. We observed two distinct subsets (Table 1: L20, L22; Supplementary Table S2, Fig 2, Supplementary Fig 1 (tissue-specific dotplot)) with a significant similarity to various *Drosophila* muscle cell types, each with a strong enrichment in different myosin (MYH, MYL) and actin (ACT) paralogs. Despite expression of several tissue-specific markers, neither of the two myocyte type could be identified to known *Drosophila* muscle types: both showed significant similarity to a variety of FCA cell types, from cardiomyocytes to skeletal muscles to visceral muscle of the midgut (Supplementary Table S2). Neither of these cell types showed any association to either SC head vs. body comparison or bulk RNAseq data, indicating that both types are present in a variety of body parts and tissues. One of these cell types (L22) can be tentatively identified as striated muscles cells, in which myosins were co-expressed with 3 different homologs of titin (TTN), numerous homologs of collagens, a homolog of troponin (TNNT3-UP), as well as invertebrate-specific muscle-specific proteins such as kettin (SLS) and paramyosin (PRM; Hooper & Thuma 2005). Additionally, ZNF593/WUPA (wings-upA) - a cytoskeletal protein of the troponin complex of the muscle thin filament - was overexpressed in this cluster. The other cell type (L20), expressing a different set of myosins, also showed elevated expression of a homolog of heart-type fatty acid binding protein FABP3 / FABP known to be associated with cardiac myocytes in mammals, and to be expressed, among other tissues, in the heart in *Drosophila.* Interestingly, the two top markers of this cell types were orthologs of *Drosophila* and human RNA endo- (*RNASET2 / RNASEX25*) and RNA exo-nucleases (*REXO5*). The significance of this observation is not clear.

#### Reproductive tissues

*Daphnia* reproductive system consists of ovaries in females (producing either parthenogenetic diploid or meiotic haploid eggs) and testes in males. Both organs are located dorsally on either side of the gut and extend from the middle section of thorax to the proximal section of the abdomen. We were able to identify one cell type unambiguously enriched in female reproductive tissues markers (L19) and, in male-derived samples, one male reproductive tissues-enriched cluster (DM2021_v1.L17), by matching to FCA markers (Table 1, Supplementary Table S2, Fig. 2, Supplementary Fig 1. Not surprisingly, these cell types were overwhelmingly over-represented in female-vs. male-derived libraries and vise versa, respectively. Comparison of markers of the female reproductive tissue cell type to those reported by Rust et al. (2020) revealed a lower degree of overlap between the most cluster-specific markers shared between *Daphnia* and *Drosophila* reproductive tissues then between neurons or myocytes of these species. Not a single among top 20 markers of cluster L19 is shared with *Drosophila* ovary-specific markers listed by Rust et al 2020. Markers of cluster 19 that are shared by these two lists, while still undoubtedly cell type-specific, showed much weaker enrichment in both. There included an ovary-specific paralogs (out of at least three in *Daphnia)* of beta-tubulin (*TUBB4B / BETATUB56D*), *penduline (KPNA2 / PEN), tet (* TET1 / TET), and cadherin (*PCDH15 / CAD99C*). Additionally, this cell type was characterized by several *Daphnia* homologs of *Drosophila* genes not listed as ovary markers, but with known ovary-related functionality and localization, such as *yolkless (LRP4 / YL*) critical for yolk deposit into oocytes, *peroxinectin-like* (PXDN / PXT) and *curly su (PXDN_CYSU).* Finally, as expected with the consideration that *Daphnia* ovaries remain mitotically active throughout adult life (Zaffagnini 1987), this cell type overexpressed several known mitosis markers, including two homologs of cyclin B *(CCNB1 / CYCB)* and *aurora A (AURKA / AURA*). Additionally, nearly all of the abovementioned markers (*LRP4 / YL*, PXDN / PXT, *PXDN_CYSU, CCNB1 / CYCB)* were shared between top markers of this cell type and the top ovary-expressed genes in Toyota et al 2017 *Daphnia* ovaries microarray transcriptome dataset. Overall the overlap between the two datasets was highly non-random (FET, P<2E-12).

We were also able to identify one of several cell types enriched in the male-derived libraries in the Experiment 2 as testes tissues (cluster DM2021_v1.L17). It was primarily characterized by high expression of homologs of mammalian *Kif11* and other Kif genes from kinesin superfamily with known or presumed role in spermatogenesis (Hara-Yokoyama et al 2019; *KIF18A / KLP67A, KIF23 / PAV, KIF11 / KLP61F*), with all

*Drosophila* counterparts showing specific expression in male germline or testes. Other highly specific markers included *Daphnia* homologs of *Drosophila* male reproductive system markers *sticky (CIT / STI*) and *kraken (* SERHL2 / KRAKEN).

#### Digestive system

*Daphnia* digestive system consists of 4 components: ectoderm-derived, cuticule-inlaid foregut and hindgut, and endoderm-derived midgut, which in turn consists of the gut itself and paired cul-de-sac appendages called hepatic cecae located in the head of the organism (Smirnov 2017). Only a single cell type has been identified as midgut cells (Table 1: L12). It shows a moderate marker overlap with *Drosophila* adult midgut and a very slight enrichment of markers associated with bulk RNAseq data from midgut. The most reliable markers of this cell type that overlap with markers of various *Drosophila* midgut cell types reported by Hung et al 2020, included several homologs of trypsins, chymotrypsin-like elastases, and other proteinases (CTRL / CG10472, MEP1B-CG15254, CELA2A / CG10472, OVCH1 / CG31267, PRSS2 / ALPHATRY, TMPRSS11D / LAMBDATRY and others), as well as an amylase (AMY1A / AMY), a paralog of APOD / NLAZ, and a paralog of FABP both different from the ones expressed, respectively, in neurons and muscle cells (see above).

Additionally, two cells types (L27 and L35) could be somewhat ambiguously identified to foregut/hindgut tissues (Table 1), with top FCA cell types being the adult esophagus and, perhaps due to ectodermal/cuticular origin, the adult tracheal cell. The ambiguity in the top five best FCA matches (Supplementary Table S2), in both cases suggesting non-random marker overlap with various eye-related tissues (pigment, cone, photoreceptor cells). Again, ectodermal/ cuticular nature of these cell types in *Drosophila* may be the source of this ambiguity.

#### Fat body and hemocytes

Fat body occurs in Daphnia in the thorax and abdomen stretching along the gut and identifiable by lipid vesicles. It has been shown to be the location of hemoglobin synthesis and hypothesized to be the tissue that generates hemocytes (Smirnov 2017). Using hemoglobins (*CYGB / GLOB1, CYGB / CG44174*) as published in situ hybridization-based marker (Goldmann et al. 1999), we identified cell type L5 as being either fat body or epipodite epithelium cells, both possibilities consistent with a significant enrichment of this cell type in the body vs. head-derived libraries (Table 1). However, matching to FCA markers revealed a more complex pattern. Top match was the pericerebral adult fat mass, not fat body, which was the 5th most significant match. Interestingly, the 2nd and 3rd highly non-random matches were *Drosophila* hemocyte and crystal cells (Supplementary Table S2). This pattern repeated with the other two cell types (L23, L31): the 3 top matches were, in different order, the pericerebral adult fat mass, hemocyte and crystal cells. Neither of the three cell types show any enrichment in the bulk RNAseq data for hemocytes (Table 1). We therefore hypothesized that these three cell types represent hemocyte- and hemoglobin-producing adipose tissue (possibly corresponding to abdominal and epipodite fat body or epipodite epithelium) and non-circulating, possibly immature, hemocytes produced in these tissues.

Conversely, cluster L15, for which not a single match to FCA markers reached the FDA<0.01 cut-off (and those that were close to that showed no consistency, Supplementary Table S2), demonstrated a highly significant marker overlap with the bulk hemocytes transcripts (Table 1). Few genes in this cluster have identifiable *Drosophila* orthologs and majority of markers shared by this cluster and bulk RNAseq data from hemocytes are *Daphnia-* unique. Consequently, there was no overlap with the markers of circulating *Drosophila* plasmatocytes of crystal cells as reported by Tattikota et al. 2020. We therefore hypothesized that this cell type represents circulating hemocytes and that their transcriptional profile diverged from that of *Drosophila* blood cells beyond recognition by examining the lists of shared markers. A striking feature of this cluster is that those few markers that do have *Drosophila* orthologs are orthologs of cuticular proteins (*CPR49AG* and others).

The presumed fat body cells, besides enrichment in homologs of FCA markers, shared a number of markers with *Drosophila* fat body cells (Croset et 2018), including collagen (*COL4A5 / COL4A1*) and several homologs of vitellogenin-like lipoprotein (*CV-D*). Two other SOD-fused vitellogenins, homologs of *Drosophila SOD1* as well as a homolog of *Drosophila* hemocyte marker glycoprotein *papilin* (*PAPLN / PPN*) were among the most specific markers of one or more of these cell types. The overlap between markers of these cell types and those reported by (Cho et al. 2020) include, besides the aforementioned hemoglobin homologs, laminins (*LAMB1 / LANB1* and *LAMC1 / LANB2*), and yet another paralog of *neural lazarillo* (*APOD / NLAZ*).

Additionally, the identity of one cell type (L26) showing ambiguous enrichment to neurons, glial cells, and follicle cells, has been hypothesized based on gene ontology of top several markers. Among them, several homologs of *Drosophila* collagen *COL41a* indicate the role of this cell type in basement membrane formation (Reinhardt et al 2023). Homologs of *yolkless (YL;* a different paralog from the ovaries marker, see Table 1) and *SPARC* point to ovaries, while *SPARC* is also expressed in hemocytes in *Drosophila.* On the other hand, *TEP2* and *FAT-SPONDIN* are markers of fat body, the latter also being a marker of hemocytes. This is consistent with the transcriptional profile of “companion plasmatocytes” (Van De Bor et al. 2015), a subpopulations of hemocytes in *Drosophila* with the function of formation of the ovarian germline stem cell niche basement membrane. We hypothesize that this functionality and cell type may be conserved across *Pancrustacea*.

#### Epithelial cells

None of the presumed cuticle/epithelial cell clusters were identified by matching to FCA, as they produced inconsistent enrichment results (Table 1). However there were three cell types detected that were characterized with several top cell type-specific markers being homologs of cuticle proteins. Five out of ten top markers of one of these cell types (L21) were crustacean-specific cuticle proteins with no or poor homology to any putative *Drosophila* orthologs. This cell type also specifically expressed a muscle cell marker *MYOT / ZORMIN*. Two other cell types identified as cuticle/epithelium cells (L25 and L34) were characterized by highly specific expression of largely non-overlapping sets of various homologs of *Drosophila* cuticle proteins (*CPR’s).* L25 expressed orthologs of *Drosophila* cuticular proteins *CPR49AA* and *CPR49AE*, not expressed in L34, which, in contrast, expressed orthologs of *CPR49AG.* Interestingly, the *Daphnia* homologs of *CPR49AA, CPR49AE*, and *CPR49AG,* which are linked in *Drosophila* (all mapping to the cytological chromosomal region 49A), also show synteny in *Daphnia*, all located on scaffold 000036F (DM3 assembly), in the order implying that nearby paralog sharing cell type specificity. Additionally, L25 differed from L34 by expressing a GMP synthase (ortholog of Drosophila BUR), and two linked paralogs with homology to neurotrophins.

### Cell type differentiation among paralogs

The analysis of cell specificity of expression of members of expanded gene families tested the hypothesis that specialization of paralogs is concordant with their sequence divergence. As expected, cell type differentiation was observed between distantly related paralogs, including orthologs of the same Drosophila genes, but not among cell type specific markers that are recent, Daphnia-specific duplicates (Fig. 3, Supplementary Table S4). Only a single pair of uncharacterized Daphnia-specific paralogs with amino acid identity over 80% showed high specificity to different cell types and those cell types were clusters L18 and L19, which are closely related and may be artifacts of clusterization (Fig. 1). In contrast, several examples of deeply differentiated paralogs being top markers of different cell types were identified (Fig.4).

**Figure 3.**
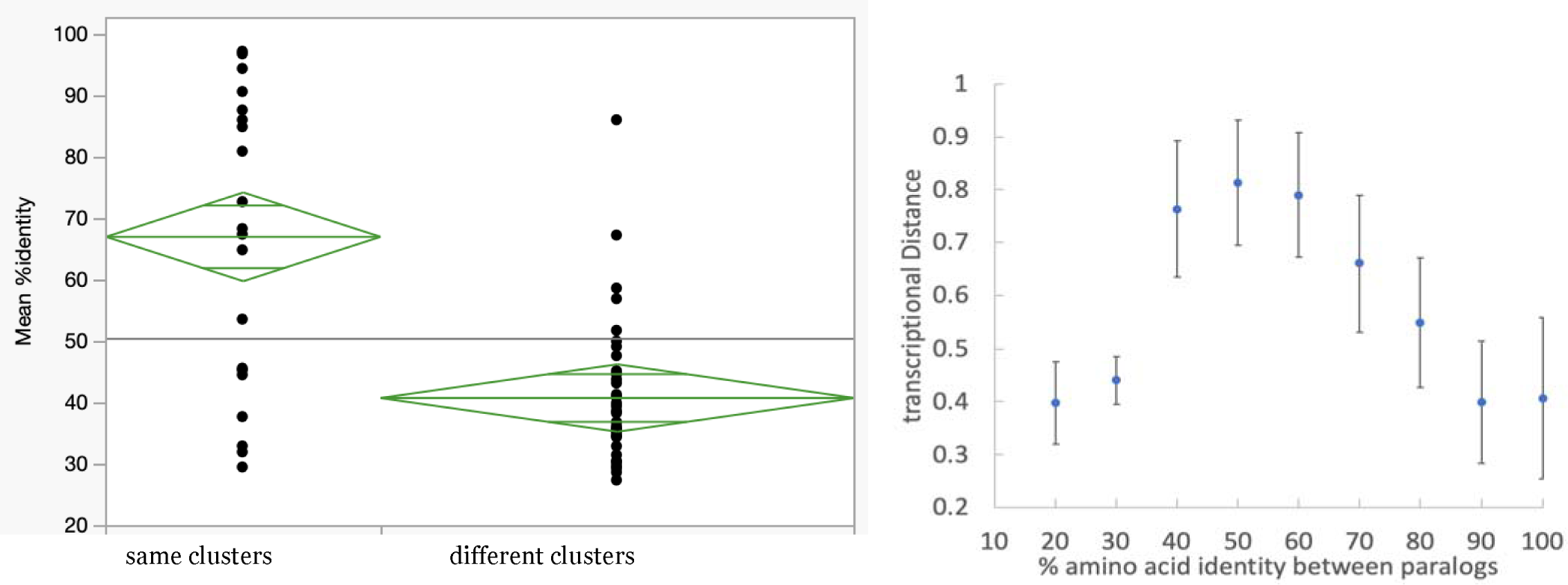
Correlations between amino acid identity and transcriptional specificity within pairs of paralogs. A. Percent identity between pairs of paralogs among top 5 most enriched markers in various cell types. The height and the width of the diamonds reflect 95% confidence interval and sample size, respectively. Between groups t-test P<0.0001. B. All paralog pairs in which both members are within top 1000 expressed genes in at least one of the cell types analyzed. Transcriptional distance calculated as euclidean distance in the space of normalized cluster-specific expression enrichment values (Z scores). Vertical bars show 99% CIs.

**Figure 4.**
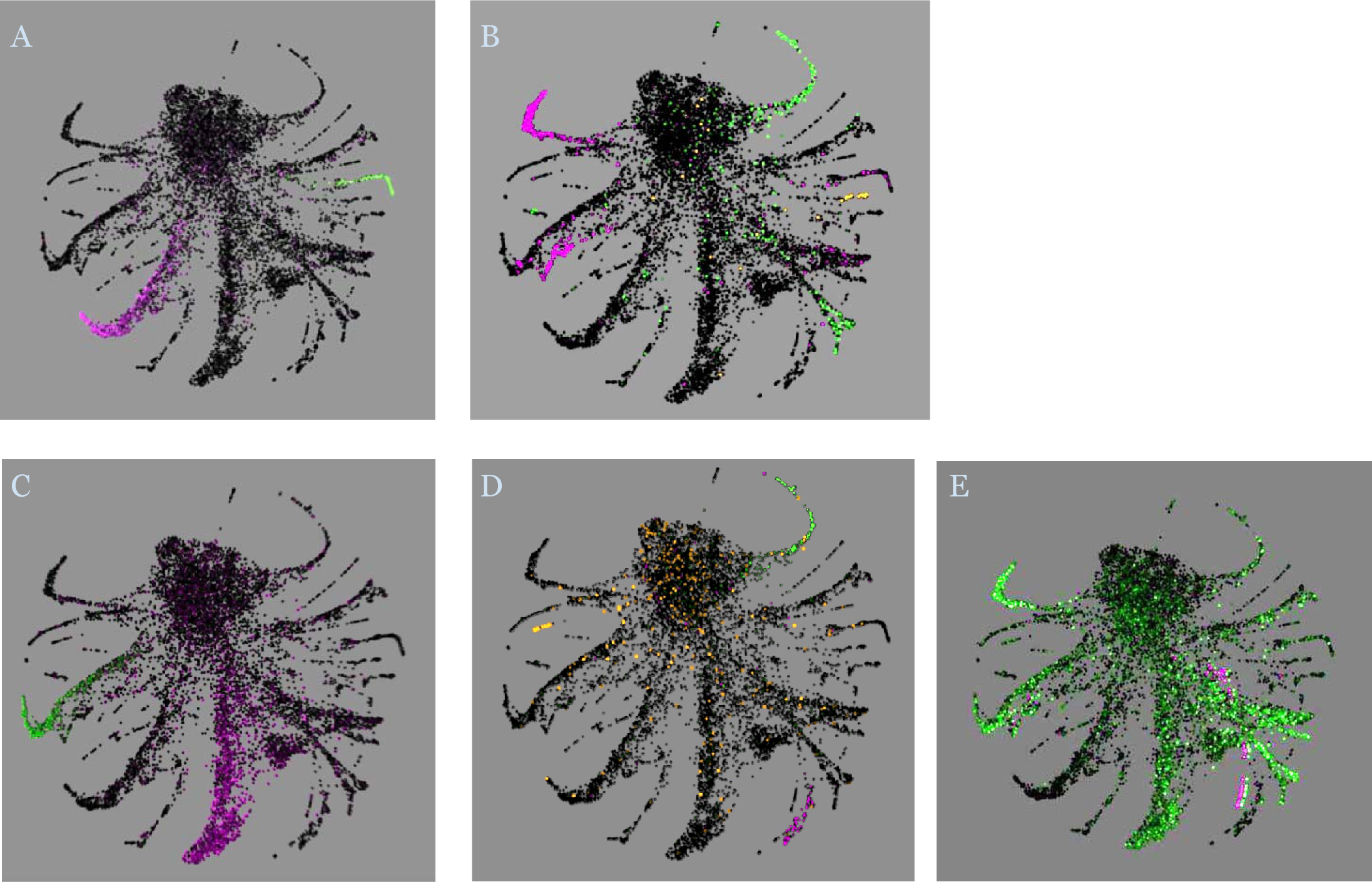
Paralog families with evidence of cell type specificity, each paralog’s expression identified as one color. A and B: orthologs of Drosophila genes yolkless (A) expressed in ovaries (purple) and fat body (green); and neural lazarillo (B), expressed in the midgut and myocytes (green), a different fat body cell type and an unidentified cluster (purple) and another unidentified cluster (orange). C - E: Daphnia paralogs identified by blast search: peroxinectin-like paralogs (C) expressed in two different neuron types, and chymotrypsin-like paralogs (D) expressed in the midgut (green), Foregut / hindgut (purple) and an unidentified cluster (orange) and ferritin heavy chain paralogs one expressed ubiquitously (green) and the other confined to an unidentified cluster (purple; cells labeled purple offset horizontally to show the other paralog’s expression in the same cell type).

The inclusion of all paralog pairs into the analysis revealed a non-monotonous relationship between transcriptional distance (a measure of specialization with respect to Louvain clusters) and percent identity (measure of homology between paralogous proteins), indicating that recent duplications share a lot of transcriptional similarity (Fig. 4B). Paralogs with 20-30 percent amino acid identity had a low transcriptional distance, which peaked at around 40% and steadily decreased at higher identity values. A closer look at 11 pairs of paralogs with identity >80% and transcriptional distance outside of the 99% confidence interval of a second degree polynomial regression of transcriptional distance over percent identity revealed several pairs in which one of the paralogs had ubiquitous expression while the other was highly expressed in a particular cell type. These include 3 pairs of cuticle proteins (data not shown) and one pair of paralogs encoding heavy chain ferritin (Fig. 4E).

## Discussion

Evolutionary divergence between branchiopods and insects is sufficient for many cell types to be lost or gained and to make it difficult to recognize some of those that are shared. This divergence exists on several levels: gene gains and losses, sequence divergence, cell types gains and losses, and transcriptional divergence between shared cell types. The former two types of divergence make orthologization difficult, the latter two result in limits to cell type matching using those genes that do show clear homology. As expected, we were able to recognize several conserved cell types, including neurons and myocytes, but none of Daphnia gas exchange or excretory tissues that are likely to be lineage-specific (see below). Likewise, there was little evidence that Drosophila trachea- or Malpigian tubules-related cell types had any match among Daphnia cell clusters. Even for conserved tissues such as CNS, particular morphological features may have evolved in insects and crustaceans in parallel (Strausfeld & Olea-Rowe 2021) and therefore may lack specific transcriptome similarity. So even for conserved cell types we were able to recognize general functionality (e.g., “neurons” or “myocytes”), but not specific cell types, even in cases when there is no doubt that such specific cell types exist in both species. On the other hand, transcriptional similarity can reveal intriguing possible homologies between seemingly different cell types. For example, even such deeply divergent organisms as molluscs and platworms show, at least in larval stage, transcriptional similarities among ciliary band cells and apical organ neurons pointing to possible homology of these structures in all lophotrochozoan larvae (Piovani et al. 2023). On the other hand, even cell types shared by all metazoans such as myocytes show lineage-specific innovations.

Apart from artifacts of microfluidics, barcoding, and clusterization, several well-defined cell types we see in Daphnia that lack homologs outside of Daphnia genomes may be such innovation specific to branchiopods or perhaps, more narrowly, to cladocerans. One such potential branch specific adaptation, not observed in Drosophila, is the cell type identifiable, by comparison to bulk RNAseq data, to circulating hemocytes (L15, Table 1; Supplementary Table S2). Two top markers of this cell type are cuticular proteins, one with and one without a close homology in Drosophila genome, and both different from cuticular proteins expressed in putative epidermis clusters (L21, L25, and L34). High-ranking markers of this cluster shared with bulk RNAs expressed in hemocytes include several linked paralogs with homology to Drosophila obstructor proteins, a family of proteins with functions related to organization and preservation of chitinous cuticle. In Drosophila these proteins are expressed largely in epithelial cells, but also, at a lesser rate in hemocytes. We hypothesize that in Daphnia the circulating hemocytes, in addition to their classic macrophagous activity, have a regeneration or wound healing function. The role of ‘blood clots’ (hemolymph clotting) and melanization in regeneration of carapace wounds has been studied in Daphnia nearly a century ago (Ermakov 1927; Anderson 1933). If the Daphnia circulating hemocytes were in fact expressing the melanization enzyme tyrosinase (the crustacean homolog of which is known as hemocyanin, and Drosophila homologs as prophenoloxidases, PPOs), it would have made them similar to Drosophila crystal cells that selective express PPOs in the fly (Ferjoux et al 2007). This is, however, not the case. The Daphnia homolog (KZS21022.1), that has a known transcriptional response to pathogens (Labbé & Little 2009) shows, in our data, a ubiquitous expression across all cell types. It appears, therefore, that this population of hemocytes accomplishes the structural wound healing function, depositing chitin and cuticular proteins, while melanization occurs by the action of hemolymph enzyme synthesized elsewhere. Crystal cells, with their ability to accumulate crystals of prophenoloxidases and specific melanization and encapsulation functions, are, therefore, likely to be an insect-specific cell type. An intriguing possibility is that initially hemocyanin was expressed in the circulating hemocytes as the oxygen carrier, but eventually assumed a new function, namely wound repair through melanization, due to the newly evolving phenoloxidase activity, with expression shifting from hemocytes to all tissues that may experience mechanical damage.

We also observe marker sharing indicative of ontological connection between hemocytes and fat body. A priori, we expected that fat body and hemocytes will be the easiest cell types to be identified, due to, respectively, the availability of hemoglobin as a marker of fat cells (Goldmann et al. 1999) and the possibility of matching transcriptional profile to that of hemocytes in bulk RNAseq data. Yet, in the FCA marker overlap analysis, all three cell types identifiable as adipocytes also showed a significant degree of marker overlap with Drosophila hemocytes and crystal cells, at the same time showing no correlation with bulk RNAseq transcripts from hemocytes. This has led us to a hypothesis that fat body shares transcriptional profile with a population of non-circulating hemocytes, possibly serving as the site where they are produced. Close association between hemocytes and fat body in crustaceans has long been hypothesized (Smirnov 2017). In one of the classic studies on Daphnia hemolymph, Hardy (1892) went as far as to suggest that the fat body is formed by recruiting lipid-rich hemocytes (which are known to become attached to non-circulatory tissues, at least temporarily). It would appear that the opposite is more likely: fat body generates hemocytes.

Another possible example of long-range evolutionary relationships between cell types is the shared similarity of Drosophila esophagus and tracheal cells to one of the inferred Daphnia cell types (L27; Table 1; Supplementary Table S2). If the less strict definition of FCA markers is used this cluster is most significantly similar to tracheal cells, with esophagus, crop and hindgut among the top 5 hits. Under the more stringent definition of FCA markers (logFC>2) tracheal cells are the 6th close match. Esophagus, foregut, crop and hindgut are all of ectodermal origin in both crustaceans and insects, essentially being cuticular invaginations into the digestive tract. We believe that this shared similarity has to do with the origin of insect trachea from similar invaginations of cuticle (Harrison 2015). A hard to interpret fact is that regardless of FCA marker stringency cut-off, one other Drosophila cell type sharing similarity to this cluster is the eye photoreceptor cells (Supplementary Table S2).

One other question we may ask is about transcriptional affinities of Drosophila organs and cell types that a priori do not have an immediate counterpart in Daphnia, for example Malpighian tubules. Tube-shape excretory organs evolved in arthropods three times independently (in insects, spiders and myriapods) each time likely to be associated with transition from aquatic to terrestrial habitats, with the primary function being the secretion of primary urine into the hindgut for water reabsorption (Benoit et al 2023). In Drosophila, Malpighian tubules originate at the midgut-hindgut boundary and, in their final form, are of dual ectoderm/mesoderm origin, some cells originating from hindgut ectoderm, some from caudal mesoderm (Jung et al 2005). No similar anatomy exists in Daphnia. The excretory organ (variably known as shell gland, maxillary gland and nephridium capsule) is located in the anterior portion of the body and does not open into the digestive tract (Smirnov 2017). It is therefore not surprising that adult fly Malpighian tubules do not show any similarity to any well-supported Daphnia cluster, except L6 which, in turn, ambiguously shares similarity with the fly midgut cells and ovaries (Supplementary Table S2). No trace of transcriptional similarity detected with any of the clusters identified as possible hindgut. On the other hand, two several specific Malpighian tubules cell types share several markers with Daphnia clusters we identified as fat body/hemocytes (L5 and L23+L31, Table 1), namely the bar-shaped cell of initial segment and the stellate cell of main segment (all FDRs for markers overlap <0.001). This may be an indication of retained transcriptional profile similarity to the mesodermal component of Malpighian tubules in Drosophila.

Another example of how transcriptional repertoire differs among cell types in the two species is offered by the photoreceptors. A small cluster of cells in Daphnia readily identifiable as photoreceptors (L36) by highly specific expression of opsins has the strongest similarity to a number of Drosophila neuron types, but not specifically to Drosophila photoreceptors, opsins expression notwithstanding. “Photoreceptor-like” and ocellus retinula cells are the closest matches, ranking 43th and 46th, respectively, with the actual Drosophila photoreceptors ranking 61st. On the other hand, for the ocellus retinula cells this Daphnia cluster is the second closest match among all Daphnia cell types, trailing only L13 (also identifiable as a neuronal cell type). On the other hand, several Daphnia clusters showing ambiguous affinities to Drosophila cell types share markers with photoreceptors (Supplementary Table S2), including, notably, L25 identified as epithelial cells by expression of cuticle proteins and collagens. Markers shared by this cell type and Drosophila photoreceptors include a neuronal markers futsch and CG11155, and Centaurin gamma 1A (CenG1A), a GTPase which is a part of ecdysone signaling-dependent pathway; the latter two also showing expression in Drosophila epithelium. Thus, Daphnia cuticle may have photoreceptor-like functionality detecting light. Presence of an autonomous, light-entrainable circadian clock affecting the cuticle and relying on a photoreceptor located in the thorax is well-documented for Drosophila (Ito et al. 2008).

In conclusion, despite deep divergence in cell types present in the two organisms and their transcriptional profiles, we were able to match some of Daphnia cell types to those in the fruitfly. Perhaps not surprisingly, the most reliable match occurred in such conserved cell types as neurons, myocytes, and midgut cells, while less specialized tissues such as fat body or hemocytes showed more ambiguous matches to the fly cell types.

## Acknowledgements

We are grateful to Dieter Ebert and Peter Fields for providing the DM3 Daphnia magna genome assembly ahead of public release and to Marc Kirschner for useful discussions. This work was supported by the NIH R24OD031956 award to L.P. and Impetus foundation grant to L.Y.

## Data availability

Raw reads are available from GEO GSE267017. Dimensionality reduction, enrichment, and clusterization results in the form of SPRING plots are available at https://shorturl.at/opAIV and https://shorturl.at/oDFM7. Enrichment of differentially expressed genes by FlyCellAtlas markers and gross anatomy differential expression are available in Supplementary data.

## Contributions

LP designed and supervised the study, IK developed the protocol and carried out experimental work, LYY participated in protocol development, cell types annotation, data analysis and manuscript preparation, KP participated in manuscript preparation and data analysis.

