## Supplementary Tables and Figures for "Single Cell Transcriptome Defines Cell Type Repertoire of Adult *Daphnia magna*"

Supplementary Table S**1**. Contingency analysis of cell type count between two protocols in Experiment 1. Each cell contains observed count, expected count and cell Chi^2^. Overall likelihood ratio Chi^2^ = 42.613, P>0.24; Pearson Chi^2^ = 40.5, P>0.31.

| Louvain cluster | Enzyme treatment protocol | | Total |
| --- | --- | --- | --- |
|  | Collagenase and hyaluronidase | Collagenase alone |  |
| 0 | 686 | 318 | 1004 |
|  | 672.4775 | 331.5225 |  |
|  | 0.2719 | 0.5516 |  |
| 1 | 388 | 169 | 557 |
|  | 373.0776 | 183.9224 |  |
|  | 0.5969 | 1.2107 |  |
| 2 | 196 | 93 | 289 |
|  | 193.5717 | 95.4283 |  |
|  | 0.0305 | 0.0618 |  |
| 3 | 493 | 243 | 736 |
|  | 492.9715 | 243.0285 |  |
|  | 0 | 0 |  |
| 4 | 305 | 139 | 444 |
|  | 297.3904 | 146.6096 |  |
|  | 0.1947 | 0.395 |  |
| 5 | 279 | 107 | 386 |
|  | 258.5421 | 127.4579 |  |
|  | 1.6188 | 3.2836 |  |
| 6 | 246 | 101 | 347 |
|  | 232.42 | 114.58 |  |
|  | 0.7935 | 1.6095 |  |
| 7 | 440 | 218 | 658 |
|  | 440.7273 | 217.2727 |  |
|  | 0.0012 | 0.0024 |  |
| 8 | 319 | 174 | 493 |
|  | 330.2105 | 162.7895 |  |
|  | 0.3806 | 0.772 |  |
| 9 | 519 | 247 | 766 |
|  | 513.0655 | 252.9345 |  |
|  | 0.0686 | 0.1392 |  |
| 10 | 20 | 7 | 27 |
|  | 18.08455 | 8.915447 |  |
|  | 0.2029 | 0.4115 |  |
| 11 | 230 | 122 | 352 |
|  | 235.769 | 116.231 |  |
|  | 0.1412 | 0.2863 |  |
| Table S1 continued 12 | 147 | 94 | 241 |
|  | 161.4214 | 79.57862 |  |
|  | 1.2884 | 2.6135 |  |
| 13 | 50 | 24 | 74 |
|  | 49.56507 | 24.43493 |  |
|  | 0.0038 | 0.0077 |  |
| 14 | 434 | 248 | 682 |
|  | 456.8024 | 225.1976 |  |
|  | 1.1382 | 2.3089 |  |
| 15 | 45 | 29 | 74 |
|  | 49.56507 | 24.43493 |  |
|  | 0.4205 | 0.8529 |  |
| 16 | 101 | 53 | 154 |
|  | 103.1489 | 50.85107 |  |
|  | 0.0448 | 0.0908 |  |
| 17 | 44 | 28 | 72 |
|  | 48.22547 | 23.77453 |  |
|  | 0.3702 | 0.751 |  |
| 18 | 364 | 191 | 555 |
|  | 371.738 | 183.262 |  |
|  | 0.1611 | 0.3267 |  |
| 19 | 230 | 93 | 323 |
|  | 216.3448 | 106.6552 |  |
|  | 0.8619 | 1.7483 |  |
| 20 | 273 | 137 | 410 |
|  | 274.6173 | 135.3827 |  |
|  | 0.0095 | 0.0193 |  |
| 21 | 36 | 18 | 54 |
|  | 36.16911 | 17.83089 |  |
|  | 0.0008 | 0.0016 |  |
| 22 | 129 | 76 | 205 |
|  | 137.3086 | 67.69136 |  |
|  | 0.5028 | 1.0198 |  |
| 23 | 100 | 50 | 150 |
|  | 100.4697 | 49.53026 |  |
|  | 0.0022 | 0.0045 |  |
| 24 | 7 | 0 | 7 |
|  | 4.688588 | 2.311412 |  |
|  | 1.1395 | 2.3114 |  |
| 25 | 84 | 40 | 124 |
|  | 83.05498 | 40.94502 |  |
|  | 0.0108 | 0.0218 |  |
| 26 | 102 | 69 | 171 |
|  | 114.5355 | 56.4645 |  |
|  | 1.372 | 2.783 |  |
| 27 | 84 | 39 | 123 |
|  | 82.38519 | 40.61481 |  |
|  | 0.0317 | 0.0642 |  |
| 28 | 16 | 10 | 26 |
|  | 17.41475 | 8.585245 |  |
|  | 0.1149 | 0.2331 |  |
| 29 | 5 | 1 | 6 |
|  | 4.01879 | 1.98121 |  |
|  | 0.2396 | 0.486 |  |
| 30 | 39 | 16 | 55 |
|  | 36.8389 | 18.1611 |  |
|  | 0.1268 | 0.2572 |  |
| 31 | 79 | 48 | 127 |
|  | 85.06438 | 41.93562 |  |
|  | 0.4323 | 0.877 |  |
| 32 | 67 | 29 | 96 |
|  | 64.30063 | 31.69937 |  |
|  | 0.1133 | 0.2299 |  |
| 33 | 10 | 8 | 18 |
|  | 12.05637 | 5.943631 |  |
|  | 0.3507 | 0.7115 |  |
| 34 | 95 | 41 | 136 |
|  | 91.09256 | 44.90744 |  |
|  | 0.1676 | 0.34 |  |
| 35 | 89 | 49 | 138 |
|  | 92.43216 | 45.56784 |  |
|  | 0.1274 | 0.2585 |  |
| 36 | 19 | 9 | 28 |
|  | 18.75435 | 9.245649 |  |
|  | 0.0032 | 0.0065 |  |
| 37 | 3 | 1 | 4 |
|  | 2.679193 | 1.320807 |  |
|  | 0.0384 | 0.0779 |  |
| Total | 6773 | 3339 | 10112 |

Supplementary Table S**2**. Top 5 FCA cell types matched to Louvain clusters markers (clusters numbered as on

https://shorturl.at/opAIV, except cluster DM2021_v1.17 which is from https://shorturl.at/oDFM7.

| Louvain  cluster | Identified as | Top 5 FCA group matches | FDR p-value |
| --- | --- | --- | --- |
| 1 | Neurons | leg muscle motor neuron | 6.41E-09 |
|  |  | labral sense organ mechanosensory neuron | 1.40E-08 |
|  |  | adult ventral nervous system | 6.28E-08 |
|  |  | scolopidial neuron | 9.72E-08 |
|  |  | adult peripheral nervous system | 5.75E-07 |
| 2 | Neurons | labral sense organ mechanosensory neuron | 3.88E-08 |
|  |  | leg muscle motor neuron | 3.96E-08 |
|  |  | multidendritic neuron | 1.16E-07 |
|  |  | adult ventral nervous system | 1.44E-07 |
|  |  | scolopidial neuron | 3.16E-07 |
| 13 | Neurons | neuron of haltere | 2.71E-32 |
|  |  | photoreceptor-like | 8.17E-26 |
|  |  | leg muscle motor neuron | 1.02E-25 |
|  |  | labral sense organ mechanosensory neuron | 3.05E-24 |
|  |  | nociceptive neuron | 7.23E-24 |
| 28 | Neurons | distal medullary amacrine neuron Dm9 | 2.90E-03 |
|  |  | bitter-sensitive labellar taste bristle | 1.47E-02 |
|  |  | olfactory receptor neuron | 1.90E-02 |
|  |  | adult olfactory receptor neuron Or47a, Or56a and likely other ORN types | 2.31E-02 |
|  |  | dopaminergic PAM neuron | 2.69E-02 |
| 36 | Neurons | adult ventral nervous system | 4.46E-11 |
|  |  | leg muscle motor neuron | 6.69E-11 |
|  |  | medullary intrinsic neuron Mi1 | 3.67E-10 |
|  |  | mechanosensory neuron | 3.11E-09 |
|  |  | adult peripheral nervous system | 3.79E-09 |
| 19 | Ovary | btl-GAL4 positive female cell, cluster 3, likely to be ovary cell | 3.67E-11 |
|  |  | post-mitotic endocycling nurse cell | 1.86E-10 |
|  |  | ovary cell | 9.61E-09 |
|  |  | germ cell stage 4 and later | 1.31E-08 |
|  |  | germline cell, unknown stage | 8.91E-08 |
| DM2021_v1.17 | Testes | spermatogonium | 1.85E-34 |
|  |  | germline cell | 6.51E-32 |
|  |  | spermatocyte 0 | 7.52E-32 |
|  |  | ejaculatory bulb epithelium | 1.24E-31 |
|  |  | ovary cell | 4.09E-31 |
| 5 | Fat body / non-circulating hemocytes | pericerebral adult fat mass | 2.14E-14 |
|  |  | crystal cell | 3.27E-09 |
|  |  | hemocyte | 8.69E-09 |
|  |  | stretch follicle cell | 3.23E-07 |
|  |  | cone cell | 6.12E-07 |
| 23 | Fat body / non-circulating hemocytes | hemocyte | 3.22E-12 |
|  |  | crystal cell | 1.36E-09 |
|  |  | leg muscle motor neuron | 1.99E-09 |
|  |  | *photoreceptor-like* | 1.12E-07 |
|  |  | *stretch follicle cell* | 3.45E-07 |
| 31 | Fat body / non-circulating hemocytes | hemocyte | 4.16E-08 |
|  |  | pericerebral adult fat mass | 6.72E-08 |
|  |  | crystal cell | 3.29E-07 |
|  |  | *stretch follicle cell* | 6.35E-06 |
|  |  | *follicle cell* | 6.05E-05 |
| 15 | Hemocytes | adult midgut* | 4.66E-06 |
|  | (identified by match to bulk RNAseq data) | midgut | 1.24E-04 |
|  |  | eo support cell | 1.76E-04 |
|  |  | adult enterocyte | 2.69E-04 |
|  |  | enterocyte-like | 1.66E-03 |
| 12 | Midgut | adult tracheal cell | 2.35E-06 |
|  |  | eye photoreceptor cell | 1.88E-03 |
|  |  | *adult esophagus* | 1.99E-03 |
|  |  | crop | 3.07E-03 |
|  |  | adult hindgut | 4.46E-03 |
| 27 | Foregut / hindgut | adult esophagus | 6.47E-05 |
|  |  | *pigment cell* | 3.46E-04 |
|  |  | crop | 5.50E-04 |
|  |  | *adult olfactory receptor neuron Or67d* | 8.42E-03 |
|  |  | *cone cell* | 1.24E-02 |
| 35 | Foregut / hindgut | cardiomyocyte, working adult heart muscle (non-ostia) | 1.22E-09 |
|  |  | *columnar neuron T1* | 8.66E-09 |
|  |  | skeletal muscle of head | 6.90E-08 |
|  |  | *choriogenic main body follicle cell and corpus luteum* | 1.22E-06 |
|  |  | muscle cell | 3.98E-06 |
| 20 | Muscle | visceral muscle of the midgut | 1.68E-25 |
|  |  | adult alary muscle | 2.38E-23 |
|  |  | cardiomyocyte, working adult heart muscle (non-ostia) | 3.64E-23 |
|  |  | adult heart ventral longitudinal muscle | 5.38E-22 |
|  |  | adult heart | 9.06E-22 |
| 22 | Muscle | eye photoreceptor cell | 1.50E-02 |
|  |  | photoreceptor cell R7 | 9.26E-02 |
|  |  | adult antenna glial cell | 1.06E-01 |
|  |  | enteroendocrine cell | 1.14E-01 |
|  |  | outer photoreceptor cell | 1.50E-01 |
| 21 | Epithelium | adult alary muscle | 2.95E-07 |
|  | (identified by cuticular proteins markers) | cone cell | 1.82E-06 |
|  |  | epithelial cell | 1.86E-06 |
|  |  | crop | 7.58E-06 |
|  |  | optic-lobe-associated cortex glial cell | 2.44E-05 |
| 25 | Epithelium |  |  |
|  | (identified by cuticular proteins markers) | enterocyte of posterior adult midgut epithelium | 4.12E-09 |
|  |  | copper cell | 2.94E-07 |
|  |  | btl-GAL4 positive female cell, cluster 2, likely to be ovary cell | 3.14E-07 |
|  |  | stretch follicle cell | 6.10E-07 |
| 34 | Epithelium; (identified by cuticular proteins markers) | germline cell | 4.65E-06 |
|  |  | crop | 6.91E-03 |
|  |  | distal medullary amacrine neuron Dm10 | 7.56E-03 |
|  |  | T neuron T2 | 1.09E-02 |
|  |  | T neuron T4/T5 | 1.13E-02 |
| Unidentified clusters | | | |
| 6 | Ambiguous:  midgut/germline. | skeletal muscle of head | 1.70E-02 |
|  | Top markers: no characterized homologs | perineurial glial sheath | 2.02E-02 |
|  |  | cone cell | 1.33E-01 |
|  |  | adult glial cell | 1.66E-01 |
|  |  | follicle cell St. 9+ | 2.15E-01 |
| 10 | Ambiguous: crop/neurons/myocyte | epithelial cell | 1.71E-03 |
|  | top markers: ambiguous | hemocyte | 2.33E-03 |
|  |  | adult tracheal cell | 4.41E-03 |
|  |  | adult abdominal pericardial cell | 5.34E-03 |
|  |  | adult midgut* | 1.34E-02 |
| 16 | Ambiguous: hemocyte/epithelium/  midgut | adult pylorus | 9.86E-05 |
|  | top markers: ambiguous | adult fat body | 2.36E-04 |
|  | note: homolog of insect- and spider silk proteins | auditory sensory neuron | 6.32E-04 |
|  |  | muscle cell | 7.94E-04 |
|  |  | enterocyte of posterior adult midgut epithelium | 1.03E-03 |
| 17 | Ambiguous: fat body/posterior gut/neurons/myocyte | distal medullary amacrine neuron Dm3 | 2.32E-02 |
|  | top markers: ambiguous | proximal medullary amacrine neuron Pm4 | 4.00E-02 |
|  |  | T neuron T4/T5c-d | 4.39E-02 |
|  |  | crop | 5.72E-02 |
|  |  | adult fat body | 6.21E-02 |
| 24 | Ambiguous: fat body/  ovary/neurons/midgut | stretch follicle cell | 3.45E-07 |
|  | top markers: collagens (non-specific) | posterior terminal follicle cell ca. St. 5-8 | 7.14E-07 |
|  |  | adult fat body | 7.16E-07 |
|  |  | adult ventral nervous system | 6.86E-06 |
|  |  | central main body follicle cell ca. St. 6-8 | 7.83E-06 |
| 26 | Ambiguous: neuron/ovary/  glia/photoreceptor |  |  |
|  | top markers: ambiguous |  |  |
| 29 | Ambiguous: neurons/oviduct/ myocyte | leg muscle motor neuron | 1.08E-10 |
|  | top markers: Daphnia-specific | follicle stem cell and prefollicle cell | 2.37E-10 |
|  |  | adult brain perineurial glial cell | 6.21E-10 |
|  |  | photoreceptor-like | 2.85E-09 |
|  |  | prefollicle cell/stalk follicle cell | 1.43E-08 |
| 30 | Ambiguous: esophagus/fat body/  germline/epidermal/  midgut | Johnston organ neuron | 7.12E-39 |
|  | cuticulum proteins among top markers | mechanosensory neuron of haltere | 1.03E-37 |
|  |  | oviduct | 6.87E-34 |
|  |  | principal cell* | 8.70E-32 |
|  |  | indirect flight muscle | 3.43E-30 |
| 33 | Ambiguous: midgut / ovary | adult esophagus | 6.92E-03 |
|  | top markers: Daphnia-specific | pericerebral adult fat mass | 8.58E-03 |
|  |  | post-mitotic germ cell early 16-cell cyst | 1.01E-02 |
|  |  | tormogen cell | 1.18E-02 |
|  |  | midgut | 1.23E-02 |

Supplementary Table S**3**. Louvain clusters excluded from the analysis based on low Z-scores of top markers (clusters numbered as on https://shorturl.at/opAIV.

| Louvain  cluster | Mean(SD) of top 5 markers Z-scores | Top 5 FCA group matches | | FDR p-value |
| --- | --- | --- | --- | --- |
| 0 | 0.941 (0.167) | adult ventral nervous system | | 1.61E-25 |
|  |  | leg muscle motor neuron | | 1.09E-16 |
|  |  | adult peripheral nervous system | | 1.87E-12 |
|  |  | nociceptive neuron | | 6.92E-12 |
|  |  | Johnston organ neuron | | 1.61E-11 |
| 3 | 0.831 (0.113) | midgut | | 1.74E-52 |
|  |  | subperineurial glial cell | | 3.12E-52 |
|  |  | ovarian sheath muscle | | 2.38E-44 |
|  |  | btl-GAL4 positive female cell, cluster 3, likely to be ovary cell | | 4.15E-44 |
|  |  | germline cell, unknown stage | | 8.14E-43 |
| 4 | 0.662 (0.191) | adult ventral nervous system | | 1.19E-07 |
|  |  | adult peripheral nervous system | | 2.33E-07 |
|  |  | labral sense organ mechanosensory neuron | | 7.84E-07 |
|  |  | lamina intrinsic amacrine neuron Lai | | 1.74E-06 |
|  |  | leg muscle motor neuron | | 6.44E-06 |
| 7 | 0.901 (0.118) | cardiomyocyte, working adult heart muscle (non-ostia) | | 4.73E-08 |
|  |  | adult ventral nervous system | | 7.70E-08 |
|  |  | labral sense organ mechanosensory neuron | | 1.47E-06 |
|  |  | adult olfactory receptor neuron Gr21a/63a | | 3.61E-06 |
|  |  | adult olfactory receptor neuron acid-sensing, Ir75a/b/c, Ir64a | | 4.81E-06 |
| 8 | 1.208 (0.11) | antimicrobial peptide-producing cell | | 2.40E-07 |
|  |  | oviduct | | 2.82E-07 |
|  |  | btl-GAL4 positive female cell, likely to be ovary cell, sim+ | | 7.32E-07 |
|  |  | choriogenic main body follicle cell and corpus luteum | | 1.77E-06 |
|  |  | cone cell | | 1.97E-06 |
| 9 | 1.053 (0.11) | oviduct | | 1.11E-11 |
|  |  | btl-GAL4 positive female cell, likely to be ovary cell, sim+, H15+ | | 6.10E-09 |
|  |  | btl-GAL4 positive female cell, likely to be ovary cell, sim+ | | 5.48E-06 |
|  |  | adult Malpighian tubule stellate cell of main segment | | 7.40E-06 |
|  |  | stretch follicle cell | | 4.09E-05 |
| 11 | 0.457 (0.187) | skeletal muscle of head | | 9.35E-95 |
|  |  | principal cell* | | 1.09E-89 |
|  |  | indirect flight muscle | | 1.40E-88 |
|  |  | epidermal cell that specialized in antimicrobial response | | 1.29E-75 |
|  |  | post-mitotic endocycling nurse cell | | 1.71E-69 |
| 14 | 0.971 (0.2) | stretch follicle cell | | 3.71E-08 |
|  |  | cardiomyocyte, working adult heart muscle (non-ostia) | | 6.26E-06 |
|  |  | adult ventral nervous system | | 7.73E-06 |
|  |  | adult Malpighian tubule stellate cell of main segment | | 2.40E-05 |
|  |  | central main body follicle cell ca. St. 6-8 | | 2.47E-05 |
| 18 | 1.038 (0.082) | btl-GAL4 positive female cell, cluster 3, likely to be ovary cell | | 3.44E-14 |
|  |  | germ cell stage 4 and later | | 3.92E-12 |
|  |  | post-mitotic endocycling nurse cell | | 4.81E-12 |
|  |  | ovary cell | | 4.84E-12 |
|  |  | post-mitotic germ cell early 16-cell cyst | | 2.27E-09 |
| 32 | 1.404 (0.351) | outer photoreceptor cell | | 0.00064815 |
|  |  | sensory neuron | | 0.00260724 |
|  |  | olfactory receptor neuron | | 0.00477104 |
|  |  | adult olfactory receptor neuron Ir84a, Ir31a, Ir76a, Ir76b, Ir8a, Or35a | | 0.00556054 |
|  |  | adult Malpighian tubule principal cell of initial segment | 0.00592435 | |

Supplementary Table S**4.** List of paralogs (triangularized matrix) with >50% coverage in at least one member of the pair that are among top 5 most enriched markers in the same or different cell types.

| gene family | Paralog 1 | | | | | Mean% identity | Paralog 2 | | | | |
| --- | --- | --- | --- | --- | --- | --- | --- | --- | --- | --- | --- |
|  | Brief ID | *Drosophila* ortholog | Cluster ID | Z-score | rank |  | Brief ID | *Drosophila* ortholog | Cluster ID | Z-score | rank |
| Same cluster | | | | | | | | | | | |
| (chemo)trypsins | m000011F_s8.62 | CG10472 | 12 | 4.161 | 1 | 32.9 | m000000F_s50.21 | ALPHATRY | 12 | 3.084 | 5 |
|  | m000011F_s8.62 |  |  | 4.161 | 1 | 29.5 | m000000F_s54.58 |  |  | 3.955 | 3 |
|  | m000000F_s54.58 |  |  | 3.955 | 3 | 45.3 | m000000F_s50.21 | ALPHATRY |  | 3.084 | 5 |
| cuticle proteins | s000036F_p3.94 | CPR49AA | 34 | 4.917 | 1 | 45.6 | m000036F_s4.48 | CPR49AE | 34 | 4.906 | 2 |
|  | s000036F_p3.94 |  |  | 4.917 | 1 | 44.5 | m000066F_s3.146 | CPR49AE |  | 4.546 | 3 |
|  | m000066F_s3.146 | CPR49AE |  | 4.546 | 3 | 87.7 | m000036F_s4.48 | CPR49AE |  | 4.906 | 2 |
| opsins | m000020F_s14.34 | RH2 | 36 | 8.886 | 1 | 97.3 | m000005F_a18.41 | RH2 | 36 | 8.23 | 2 |
|  | s000031F_p1.91 | RH2 |  | 8.039 | 3 | 96.9 | m000005F_a18.41 |  |  | 8.23 | 2 |
|  | s000031F_p1.91 |  |  | 8.039 | 3 | 97.1 | m000020F_s14.34 | RH2 |  | 8.886 | 1 |
|  | s000031F_p1.91 |  |  | 8.039 | 3 | 67.5 | m000031F_s0.69 | RH2 |  | 4.798 | 5 |
|  | m000031F_s0.69 | RH2 |  | 4.798 | 5 | 67.5 | m000005F_a18.41 | RH2 |  | 8.23 | 2 |
|  | m000031F_s0.69 |  |  | 4.798 | 5 | 64.9 | m000020F_s14.34 | RH2 |  | 8.886 | 1 |
| SOD fused with vitellogenin | m000004F_a31.54 | SOD1 | 5 | 3.613 | 2 | 90.8 | m000004F_a31.49 | SOD1 | 5 | 3.661 | 1 |
| unknown 1 | m000038F_a8.149 |  | 27 | 5.133 | 5 | 31.9 | m000018F_a7.55 |  | 27 | 5.134 | 4 |
|  | m000008F_s17.30 |  | 30 | 8.753 | 1 | 72.8 | m000008F_s17.24 |  | 30 | 8.42 | 2 |
| unknown 2 | m000018F_s7.81 |  | 15 | 5.303 | 5 | 85 | m000018F_s7.64 |  | 15 | 5.341 | 4 |
| unknown 3 | m000018F_s10.5 |  | 19 | 3.017 | 1 | 86.2 | m000018F_a6.24 |  | 19 | 2.959 | 2 |
| unknown 4 | m000006F_s30.100 |  | 25 | 4.826 | 3 | 37.7 | m000006F_a30.137 |  | 25 | 4.565 | 5 |
| unknown 5 | m000018F_a7.59 |  | 32 | 1.205 | 4 | 53.6 | g000018F_p7.121 |  | 32 | 1.293 | 3 |
| unknown 6 | g000016F_p16.34 |  | 35 | 6.442 | 1 | 68.4 | a000023F_p13.43 |  | 35 | 5.532 | 5 |
|  | s000016F_p16.68 |  |  | 6.273 | 2 | 81 | a000023F_p13.43 |  |  | 5.532 | 5 |
|  |  |  |  | 6.273 | 2 | 94.5 | g000016F_p16.34 |  |  | 6.442 | 1 |
| Different clusters | | | | | | | | | | | |
| (chemo)trypsins | m000011F_s8.62 | CG10472 | 12 | 4.161 | 1 | 35.7 | a000023F_p12.7 | CG4613 | 33 | 12.411 | 4 |
|  |  |  | 12 | 4.161 | 1 | 34.4 | m000001F_a47.66 | L(2)K05911 | 37 | 22.239 | 1 |
|  | m000000F_s54.58 |  | 12 | 3.955 | 3 | 32.9 | a000023F_p12.7 | CG4613 | 33 | 12.411 | 4 |
|  | m000000F_s50.21 | ALPHATRY | 12 | 3.084 | 5 | 40 | a000023F_p12.7 | CG4613 | 33 | 12.411 | 4 |
|  | m000001F_a47.66 | L(2)  K05911 | 37 | 22.24 | 1 | 35.5 | a000023F_p12.7 | CG4613 | 33 | 12.411 | 4 |
|  |  |  | 37 | 22.24 | 1 | 36.8 | m000000F_s50.21 | ALPHATRY | 12 | 3.084 | 5 |
|  |  |  | 37 | 22.24 | 1 | 35.1 | m000000F_s54.58 |  | 12 | 3.955 | 3 |
| collagens | m000035F_s5.102 |  | 2 | 4.35 | 2 | 39.9 | m000000F_a29.50 | COL4A1 | 24 | 27.572 | 1 |
|  | s000001F_p14.25 | COL4A1 | 26 | 4.468 | 5 | 40.9 | m000000F_a29.50 | COL4A1 | 24 | 27.572 | 1 |
|  | s000001F_p14.25 | COL4A1 | 26 | 4.468 | 5 | 34.4 | m000035F_s5.102 |  | 2 | 4.35 | 2 |
| cuticle proteins | m000036F_s4.53 | CPR49AG | 25 | 4.803 | 4 | 31.5 | m000031F_s2.44 |  | 21 | 8.652 | 1 |
| cuticle proteins |  |  | 25 | 4.803 | 4 | 58.7 | m000036F_s4.42 | CPR49AG | 15 | 5.516 | 2 |
| cuticle proteins |  |  | 25 | 4.803 | 4 | 38.3 | m000036F_s4.48 | CPR49AE | 34 | 4.906 | 2 |
| cuticle proteins | m000086F_s0.47 |  | 30 | 5.943 | 5 | 51.8 | m000036F_s4.42 | CPR49AG | 15 | 5.516 | 2 |
| cuticle proteins |  |  | 30 | 5.943 | 5 | 36.1 | m000036F_s4.48 | CPR49AE | 34 | 4.906 | 2 |
| cuticle proteins |  |  | 30 | 5.943 | 5 | 49.1 | m000036F_s4.53 | CPR49AG | 25 | 4.803 | 4 |
| cuticle proteins |  |  | 30 | 5.943 | 5 | 32.9 | m000066F_s3.146 | CPR49AE | 34 | 4.546 | 3 |
| cuticle proteins | s000036F_p3.94 | CPR49AA | 34 | 4.917 | 1 | 41.3 | m000036F_s4.42 | CPR49AG | 15 | 5.516 | 2 |
| cuticle proteins |  |  | 34 | 4.917 | 1 | 43.8 | m000036F_s4.53 | CPR49AG | 25 | 4.803 | 4 |
| cuticle proteins |  |  | 34 | 4.917 | 1 | 39.4 | m000086F_s0.47 |  | 30 | 5.943 | 5 |
| cuticle proteins | m000036F_s4.48 | CPR49AE | 34 | 4.906 | 2 | 28.6 | m000036F_s4.42 | CPR49AG | 15 | 5.516 | 2 |
| cuticle proteins | m000066F_s3.146 | CPR49AE | 34 | 4.546 | 3 | 29.3 | m000021F_a15.33 |  | 27 | 5.178 | 3 |
| cuticle proteins |  |  | 34 | 4.546 | 3 | 34.6 | m000036F_s4.42 | CPR49AG | 15 | 5.516 | 2 |
| cuticle proteins |  |  | 34 | 4.546 | 3 | 40 | m000036F_s4.53 | CPR49AG | 25 | 4.803 | 4 |
| peroxinectin-like | m000017F_a22.36 |  | 9 | 0.916 | 5 | 27.3 | m000001F_s29.79 | PXT | 2 | 4.18 | 4 |
| unknown 1 | m000013F_s8.96 |  | 15 | 5.864 | 1 | 45.2 | m000008F_s17.24 |  | 30 | 8.42 | 2 |
| unknown 1 |  |  | 15 | 5.864 | 1 | 57 | m000008F_s17.30 |  | 30 | 8.753 | 1 |
| unknown 1 |  |  | 15 | 5.864 | 1 | 47.7 | m000008F_s17.31 |  | 25 | 4.923 | 2 |
| unknown 1 | m000024F_s13.68 |  | 17 | 5.163 | 2 | 38.6 | m000008F_s17.24 |  | 30 | 8.42 | 2 |
| unknown 1 |  |  | 17 | 5.163 | 2 | 35.9 | m000008F_s17.30 |  | 30 | 8.753 | 1 |
| unknown 1 |  |  | 17 | 5.163 | 2 | 43.2 | m000008F_s17.31 |  | 25 | 4.923 | 2 |
| unknown 1 |  |  | 17 | 5.163 | 2 | 30.3 | m000013F_s8.96 |  | 15 | 5.864 | 1 |
| unknown 1 | m000008F_s17.31 |  | 25 | 4.923 | 2 | 44.6 | m000008F_s17.24 |  | 30 | 8.42 | 2 |
| unknown 1 |  |  | 25 | 4.923 | 2 | 50 | m000008F_s17.30 |  | 30 | 8.753 | 1 |
| unknown 3 | m000018F_s10.5 |  | 19 | 3.017 | 1 | 86.2 | m000018F_a6.24 |  | 18 | 0.931 | 5 |
| unknown 5 | g000018F_p7.121 |  | 32 | 1.293 | 3 | 29.7 | a000018F_p8.3 |  | 16 | 3.404 | 5 |
| unknown 5 | m000018F_a7.59 |  | 32 | 1.205 | 4 | 30.5 | a000018F_p8.3 |  | 16 | 3.404 | 5 |
| unknown 7 | m000013F_a25.98 |  | 35 | 6.14 | 4 | 67.3 | m000013F_a1.94 |  | 37 | 13.055 | 5 |

Fig. S**1**. Dotplots showing top 10 markers for reproductive system (A) and neuronal (B) cell types.

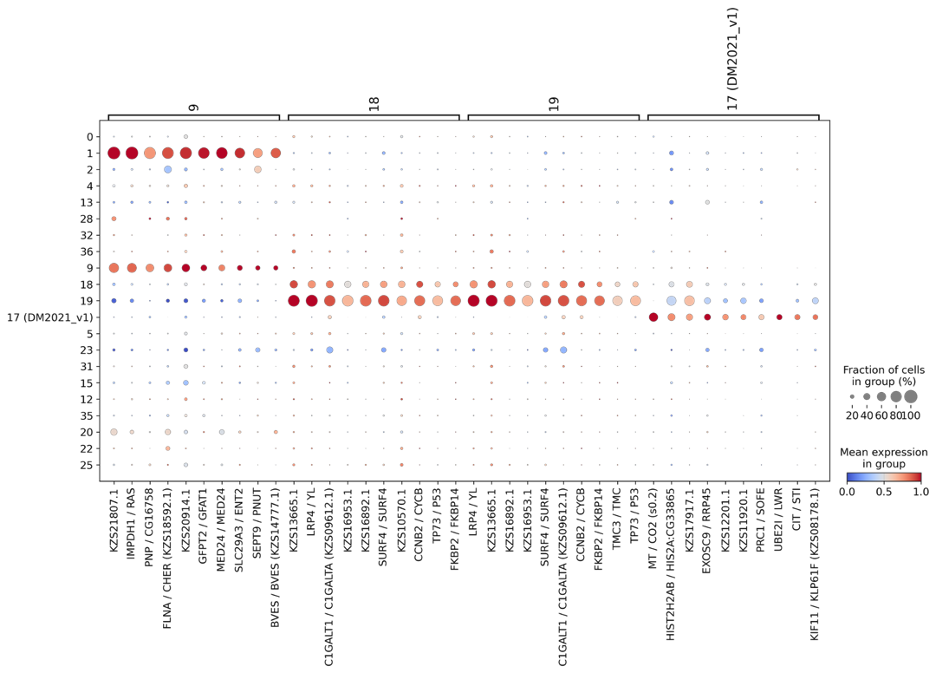

A

B

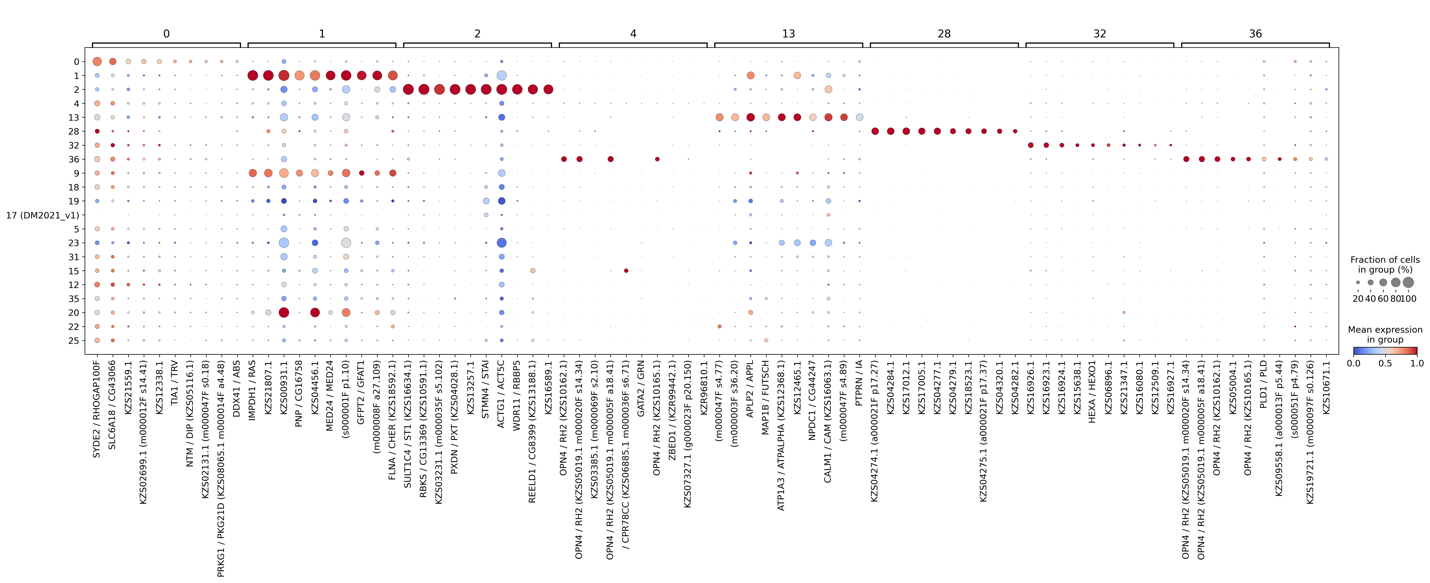
